# Systemic sterile induced-co-expression of IL-12 and IL-18 drives IFN-γ-dependent activation of microglia and recruitment of MHC-II-expressing inflammatory monocytes into the brain

**DOI:** 10.1101/2020.07.30.228098

**Authors:** Emilia A. Gaviglio, Javier M. Peralta Ramos, Daniela S. Arroyo, Claudio Bussi, Pablo Iribarren, Maria C. Rodriguez-Galan

## Abstract

The development of neuroinflammation, as well as the progression of several neurodegenerative diseases, has been associated with the activation and mobilization of the peripheral immune system due to systemic inflammation. However, the mechanism by which this occurs remains unclear. Herein, we addressed the effect of systemic, endotoxin-free induced-co-expression of IL-12 and IL-18 in the establishment of a novel cytokine-mediated model of neuroinflammation. Following peripheral hydrodynamic shear of IL-12 plus IL-18 cDNAs in C57BL/6 mice, we induced systemic and persistent level of IL-12, which in turn promoted the elevation of circulating pro-inflammatory cytokines TNF-α and IFN-γ, accompanied with splenomegaly. Moreover, even though we identified an increased gene expression of both TNF-α and IFN-γ in the brain, only TNF-α was shown to be dispensable, revealing an IFN-γ-dependent activation of microglia and the recruitment of leukocytes, particularly of highly activated inflammatory monocytes. Taken together, our results argue for a systemic cytokine-mediated establishment and development of neuroinflammation, having identified IFN-γ as a potential target for immunotherapy.

## INTRODUCTION

There is increasing evidence indicating that systemic inflammatory events can exert devastating effects in the brain due to their impact in the progression of several central nervous system (CNS) disorders, such as autoimmune and neurodegenerative diseases, in which leukocyte recruitment is a key feature (1–4).

Several experimental models, mainly associated to bacterial (5), viral (6,7) and parasitic (8,9) infections have been developed to better understand the communication between the peripheral immune system and the CNS. In this sense, lipopolisaccharide (LPS) constitutes the most commonly used stimulus to mimic systemic inflammation, due to the robust immune response exerted upon administration (2,10–12).

Our previous studies demonstrate that following a systemic LPS challenge, we induced glial reactivity together with an active type I interferon-dependent recruitment of inflammatory monocytes that were able to enhance T cell proliferation, unlike microglia (13). Besides, we have previously reported that LPS-primed inflammatory monocytes could also internalize α-synuclein, which in turn favored the dissemination of this pathogenic protein from the periphery toward the brain and spinal cord during synucleinopathies, such as Parkinson’s disease (14).

For years, interleukin (IL)-12 has been proposed as a promising candidate for cancer treatment because of its ability to elicit robust antitumor responses, not only when delivered locally, but systemically (15–18). Thus, it is currently being used in a phase 1 trial in patients with recurrent high grade glioma with favorable preliminary results (19). Moreover, accumulating data points to a central role for IL-12 in neuroinflammation (20) and Alzheimer’s disease (21,22).

Interleukin-18 is an inflammatory cytokine of the IL-1 family which has been extensively described as a key immune regulator during chronic inflammation (23), autoimmune diseases (24), infection (25) and cancer (26–28). Constitutively expressed by brain resident cells, previous studies have highlighted a crucial role for IL-18 in mediating neuroinflammation and neurodegeneration (29) upon activation of the innate immune-related inflammasome through cell-death mechanisms, such as pyroptosis (30).

Based on these results, and given that fully understanding the impact of systemic inflammation in leukocyte trafficking into the CNS is still warranted, we sought to assess the effect of peripheral sterile inflammation in the establishment of a novel cytokine-mediated model of neuroinflammation after systemic expression of IL-12 and IL-18.

## MATERIAL & METHODS

### Mice

Wild-type (WT) C57BL/6 mice were originally obtained from Escuela de Veterinaria, Universidad Nacional de la Plata. IFN-γ^-/-^ (B6.129S7-Ifngtm1Ts/J strain) and TLR4^-/-^ mice were obtained from The Jackson Laboratory, whereas IL-4^-/-^ and TNFαR1^-/-^ mice were kindly provided by Dr. Silvia Di Genaro. Between 8- and 12-week-old female mice were maintained in the specific pathogen-free barrier facilities, housed under a normal 12 h light/dark cycle with free access to pelleted food and water, at the animal facility from Facultad de Ciencias Químicas, Universidad Nacional de Córdoba. All experiments were performed in compliance with the procedures outlined in the “Guide for the Care and Use of Laboratory Animals” (NIH Publication N° 86-23, 1985). Experimental protocols were approved by the Institutional Animal Care and Use Committee (IACUC). Our animal facility obtained NIH animal welfare assurance (N° A5802-01, OLAW, NIH, US).

### Hydrodynamic injection of IL-12 and IL-18 cDNAs

Plasmids with the incorporated sequences were amplified in bacteria cultured in LB medium supplemented with ampicillin, and cDNA was purified using Endofree^®^ Plasmid Maxi Kit (Qiagen) according to manufacturer’s instructions.

The hydrodynamic gene transfer procedure was adapted according to Liu F *et al*. (31) and routinely used in our laboratory (26,27,32,33). Briefly, through this technique, the large volume and high injection rate used, forces the flow of cDNA solution into tissues directly linked to the inferior vena cava. A large portion of cDNA solution is then forced into the liver and plasmid cDNA molecules are likely transferred inside the liver cells by the hydrodynamic process during cDNA administration. In the present study, animals were injected in the tail vein with the cDNA in 1.6 ml of sterile 0.9% sodium chloride solution in < 8 sec and then separated into two groups: 1) Ctrl (11 μg of ORF empty vector control cDNA); 2) IL-12 plus IL-18 (1 μg of IL-12 cDNA, pscIL-12, p40-p35 fusion gene) plus 10 μg of IL-18 cDNA (pDEF pro-IL-18). All the expression plasmids used the human elongation 1-α promoter to drive the transcription of their respective cytokines.

### LPS challenge

Lipopolysaccharide from *Escherichia coli* 055:B5 (purified by gel-filtration chromatography) was purchased from Sigma-Aldrich and freshly dissolved in sterile saline prior to intraperitoneal (i.p.) injection. Mice were treated with either vehicle or 1.6 mg/kg of LPS for four consecutive days to induce neuroinflammation, following an injection scheme modified from Cardona *et al*. (11).

### Isolation of immune cells from mice brains

Following seven days upon initiation of the treatment, mice were weighed and deeply anesthetized with a ketamine/xylazine cocktail accordingly. Immune cells were isolated from whole brain homogenates as follows. Briefly, mice were transcardially perfused with ice-cold PBS 1x (Gibco), and brains were collected in DMEM (Gibco) supplemented with sodium pyruvate (Gibco) and a penicillin, streptomycin and glutamine cocktail (Gibco), gently mechanically disaggregated and resuspended in PBS 1x containing 3 mg/ml of collagenase D (Roche Diagnostics) plus 10 μg/ml of DNAse (Sigma-Aldrich) for an enzymatic homogenization. After a 30 min incubation, brain homogenates were filtered with 40 μm pore size cell strainers (BD Biosciences), centrifuged 8 min at 1800 r.p.m., washed with PBS 1x, and resuspended in 6 ml of 38% isotonic Percoll^®^ (GE Healthcare) before a 25 min centrifugation at 800 G without neither acceleration nor brake. Myelin and debris were discarded. Cell pellets containing total brain immune cells were collected, washed with DMEM supplemented with 10% fetal bovine serum (Gibco), and cell viability was determined by trypan blue exclusion using a Neubauer’s chamber. Finally, cells were labeled for subsequent flow cytometric analysis.

### Flow Cytometric Analysis

Surface staining of single-cell suspension of isolated brain immune cells was performed using standard protocols and analyzed on a FACSCanto II (BD Biosciences). Flow cytometric analysis was defined based on the expression of CD11b, CD45, Ly6C, CD4, and CD8 as follows: microglia, CD11b^+^ CD45^lQ^; recruited leukocytes, CD11b^+/-^ CD45^hi^; inflammatory monocytes, CD11b^+^ CD45^hi^ Ly6C^hi^ Ly6G^-^; neutrophils, CD11b^+^ CD45^hi^ Ly6G^+^ Ly6C^int^; dendritic cells, CD11b^+^ CD45^hi^ CD11c^+^; CD4 T cells, CD11b^-^ CD3^+^ CD45^hi^ CD4^+^; CD8 T cells, CD11b^-^ CD3^+^ CD45^hi^ CD8^+^; B cells, CD11b^-^ CD19^+^ CD45^hi^. Data analysis was conducted using FCS Express (De Novo Software). The following antibodies were used in the procedure: monoclonal anti-mouse CD11b APC (BioLegend, clone M1/70); CD11b Alexa Fluor^®^ 488 (BD Pharmingen, clone M1/70); CD45 APC-Cy7 (BioLegend, clone 30-F11); CD11c APC (BD Pharmingen, clone HL3); CD11c PE (BD Pharmingen, clone HL3); Ly6C PE-Cy7 (BD Pharmingen, clone AL-21); Ly6G PE (BD Pharmingen, clone 1A8); CD3 PE-Cy5 (BD Pharmingen, clone 17A2); CD4 FITC (BD Pharmingen, clone RM4-4); CD8a PE (BD Pharmingen, clone 53-6.7); CD19 PE-Cy7 (BD Pharmingen, clone 1D3); I-A/I-E Alexa Fluor^®^ 647 (BioLegend, clone M5/114.15.2) or isotype control antibodies (BD Pharmingen, APC, clone R35-95; Alexa Fluor 488^®^, clone A95-1; FITC, clone A95-1; APC-Cy7, clone A95-1; PE, clone A95-1; PE-Cy5, clone A95-1; PE-Cy7, clone G155-178). Multiparametric gating analysis strategy was performed as previously described (13,14).

### ELISA assay

Serum samples obtained at day 0 and at different time points following initiation of the treatment were tested for murine IL-12 (p70, eBioscience), TNF-α (eBioscience), IFN-γ (BD Pharmingen) and IL-10 (BD Pharmingen) according to manufacturer’s instructions.

### Splenomegaly assessment

Seven days after initiation of the treatment, mice were weighed and deeply anesthetized with a ketamine/xylazine cocktail accordingly. Spleens were then excised, weighed and photographed. In order to normalize the weight of the spleen according to each individual mouse body weight, an index was calculated as follows: (spleen weight/body weight)/average of spleen weight derived from Ctrl mice.

### Reverse transcription of mRNA and quantification by real-time PCR

Brain homogenates were resuspended in TRIzol^®^ (Invitrogen), then RNA was extracted according to manufacturer’s instructions and stored at −80°C. Total RNA was quantified using a Synergy HT spectrophotometer (BioTek) and 1 μg was treated with DNAse (Sigma-Aldrich) and reverse-transcribed using the High-Capacity cDNA Reverse Transcription Kit (Applied Biosystems). Real-time PCR was performed in a StepOnePlus™ real-time PCR system (Applied Biosystems) using SYBR^®^ Green real-time PCR master mix (Applied Biosystems), and relative quantification (RQ) was calculated by using StepOne™ software V2.2.2, based on the equation RQ= 2^-ΔΔCt^, where Ct is the threshold cycle to detect fluorescence. Ct data were normalized to the internal standard HPRT1. Primer sequences were as follows (5’-3’): TNF-α, sense: AGC CGA TGG GTT GTA CCT TGT CTA, anti-sense: TGA GAT AGC AAA TCG GCT GAC GGT; IFN-γ, sense: GGA ACT GGC AAA AGG ATG, anti-sense: GAT GGC CTG ATT GTC TTT CAA GA; HPRT1, sense: TCA GTC AAC GGG GGA CAT AAA, anti-sense: GGG GCT GTA CTG CTT AAC CAG.

### Fluorescence confocal microscopy

Following seven days upon the treatment, mice were weighed and deeply anesthetized with a ketamine/xylazine cocktail accordingly. Animals were transcardially perfused once with ice-cold PBS 1x (Gibco) and then with 4% paraformaldehyde (PFA). Brains were collected in 4% PFA for an additional 24 h postfixation and incubated in 20% sucrose for another 24 h.

For immunofluorescence assay, brains were embedded in Tissue-Tek^®^ optimal cutting temperature compound (Sakura), cut into 10 μm sections of thickness using a Shandon Cryotome E cryostat (Thermo Scientific), and mounted on Starfrost^®^ adhesive slides (Knittel Glass). Sections were rehydrated with blocking buffer (10% BSA, 0.3% Triton in TBS), rinsed with TBS (Gibco), and incubated overnight at 4°C with the corresponding dilutions of the antibodies CD45 (BioLegend, clone 30-F11) and CD31 (Santa Cruz, clone M-20) in blocking buffer. After several rinses, sections were incubated with Alexa Fluor 488 (Molecular Probes) or Alexa Fluor 546 (Molecular Probes) antibodies and counterstained with DAPI. Slides were analyzed under a FV1000 laser scanning confocal fluorescence microscope (Olympus). Quantification of microglia numbers and CD45 integrated density area was performed using ImageJ software (NIH, USA).

### Statistical analysis

Results are expressed as mean ± s.e.m. All statistical analyses were performed using Prism^®^ 7.0 (GraphPad Software). Means between two groups was compared with Student *t*-test or two-way analysis of variance followed by a Bonferroni *post-hoc* test for more than two groups. Statistical significance levels were set as follows: * if p < 0.05, ** if p < 0.01, and *** if p < 0.001.

## RESULTS

### In vivo treatment with IL-12 + IL-18 cDNAs increase circulating pro-inflammatory cytokine levels and induce splenomegaly

In both human sepsis and murine experimental models based on LPS administration, increased levels of cytokines can be detected in peripheral blood (34,35). We have previously demonstrated that hydrodynamic shear of IL-12 and IL-18 cDNAs induced systemic expression of several Th1 and Th2 type cytokines (26).

In the present study, we have used a lower dose of IL-12 in order to mimic a physiological model of sterile peripheral inflammation. Therefore, we used a single challenge of IL-12 (1 μg) plus IL-18 (10 μg) cDNAs to promote the elevation of circulating pro-inflammatory cytokines. At these doses, we noticed that administration of IL-12 plus IL-18 ramped up the level of tumor necrosis factor (TNF)-α starting from day 2 and remaining high until day 6 (**Fig. 1A**). Instead, IFN-γ rose from the first day, reaching a peak on day 2 and subsequently declining through day 3 to day 6, although remaining higher than the levels observed in control mice (**Fig. 1A**). Interleukin-10 sera levels showed a kinetic similar to IFN-γ, reaching a peak at day 2 with values forty-fold lower, that gradually decreased but remained significantly higher in comparison to their control littermates (**Fig. 1A**). Interestingly, we consistently found that this pro/anti-inflammatory cytokine profile was associated with a dramatic splenomegaly observed on day 7 upon the hydrodynamic challenge of IL-12 plus IL-18 cDNAs (**Fig. 1B**). For this reason, we considered the presence of splenomegaly in IL-12 and IL-18-treated mice as an indication of an effective hydrodynamic shear.

**Fig. 1.**
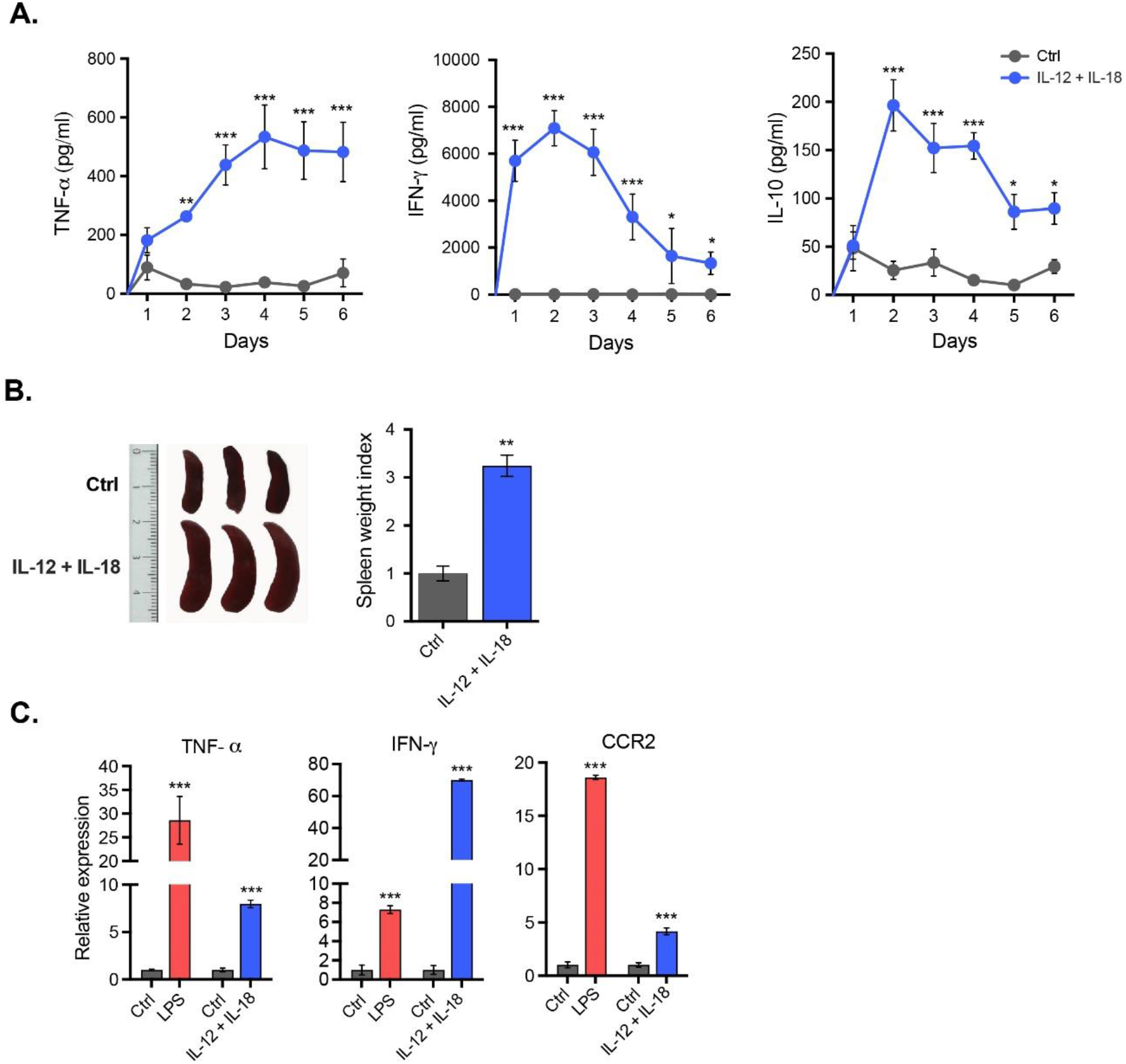
Peripheral induced-co-expression of IL-12 plus IL-18 increase circulating pro-inflammatory cytokine levels and induce splenomegaly. C57BL/6 WT mice were hydrodynamically injected with either empty vector control cDNA (Ctrl), or IL-12 plus IL-18 cDNAs (IL-12 + IL-18). **(A)** Serum samples obtained at day 0 and at different time points following initiation of the treatment were tested for TNF-α and IFN-γ. Results are representative of at least three independent experiments (n= 6-13 samples per time point and group). **(B)** Seven days after initiation of the treatment mice were euthanized, and spleens excised for assessment of their relative weight and size. Results are representative of at least three independent experiments (n= 2-3 mice per group). **(C)** Gene expression analysis by real-time qPCR of TNF-α and IFN-γ in total bulk brain. Relative quantification (RQ) was calculated based on the equation RQ= 2^-ΔΔCt^, where Ct is the threshold cycle to detect fluorescence. Ct data were normalized to the internal standard HPRT1. Results are representative of at least three independent experiments (n= 3-12 mice per group). Data are expressed as mean ± s.e.m. Means between groups was compared with Student *t*-test. Statistical significance levels were set as follows: * if p < 0.05, ** if p < 0.01, and *** if p < 0.001.

Interestingly, systemic induced-co-expression of IL-12 plus IL-18 cDNAs was able to exert a potent mRNA expression of both TNF-α and IFN-γ, as well as of CCR2 chemokine receptor in the brain, seven days after the treatment **(Fig. 1C)**. Remarkably, IFN-γ mRNA gene expression was ten-fold higher than the observed in endotoxemic mice, upon a serial LPS injection regime to induce neuroinflammation **(Fig. 1C)**.

These results clearly demonstrate that a single challenge of IL-12 and IL-18 cDNAs induce an increase in the circulating levels of pro-inflammatory cytokines, which result in a striking splenomegaly and robust mRNA expression of pro-inflammatory cytokines in the brain.

### Systemic sterile induced-co-expression of IL-12 and IL-18 induce microglia activation and favors leukocyte recruitment into the brain

Immunological surveillance of the CNS has been shown to be dynamic, specific, and tightly regulated (36). During neuroinflammation or under neurodegenerative conditions, the blood–brain barrier (BBB) might get breached, enabling the entry of peripheral immune cells through the choroid plexus (37), and subsequently into the brain parenchyma. Brain-resident microglia then encounter immune cells that have been primed in the periphery, leading to the establishment of an interplay that could worsen the outcome of the inflammatory process (38).

Given the elevation in the gene expression of TNF-α and IFN-γ in the brain upon systemic delivery of IL-12 and IL-18 cDNAs, we wondered how this pro-inflammatory microenvironment could affect both microglia activation state, and the immune response deployed from the periphery. By leveraging multiparametric flow cytometry, we found that hydrodynamic injection of IL-12 plus IL-18 was not able to modify the absolute number of CD45^lo^ CD11b^+^ microglia in the brain (**Figs. 2A and 2B**), but to induce a marked activation, as shown by the upregulation of major histocompatibility class-II (MHC-II) (**Fig. 3**).

**Fig. 2.**
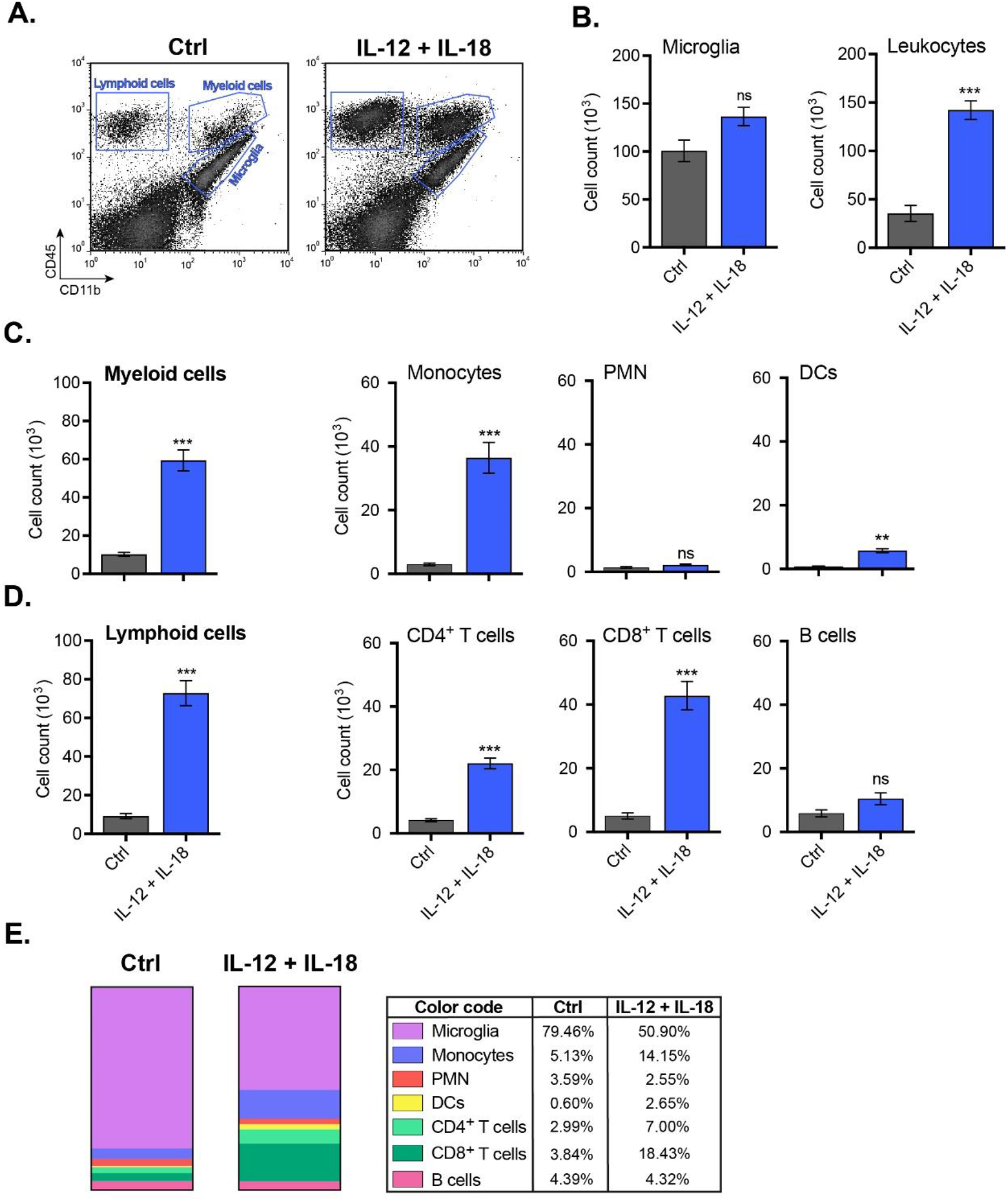
Systemic hydrodynamic shear of IL-12 plus IL-18 results in remodeling of the brain immune landscape. C57BL/6 WT mice were hydrodynamically injected with either empty vector control cDNA (Ctrl), or IL-12 plus IL-18 cDNAs (IL-12 + IL-18). Seven days after initiation of the treatment mice were euthanized, and immune cells were isolated from whole brain and stained for subsequent flow cytometric analysis. **(A)** Representative CD45 vs. CD11 b flow cytometric density-plots illustrating the gating analysis strategy employed. **(B)** Absolute number of CD11b^+^ CD45^lo^ microglia, CD11b^+/-^ CD45^hi^ recruited leukocytes, **(C)** CD11b^+^ CD45^hi^ recruited myeloid cells, CD11b^+^ CD45^hi^ Ly6C^hi^ Ly6G^-^ inflammatory monocytes, CD11b^+^ CD45^hi^ Ly6G^hi^ Ly6C^int^ PMN, CD11b^+^ CD45^hi^ CD11c^+^ DCs, **(D)** CD11b^-^ CD45^hi^ recruited lymphoid cells, CD11b^-^ CD45^hi^ CD3^+^ CD4^+^ T cells, CD11b^-^ CD45^hi^ CD3^+^ CD8^+^ T cells, and CD11b^-^ CD45^hi^ CD19^+^ B cells were assessed by flow cytometry. **(E)** Representative stacked bar chart, comparing Ctrl vs. IL-12 + IL-18 frequency of resident and peripheral CD45^+^ populations previously analyzed by flow cytometry. Results are representative of at least three independent experiments (n= 3-12 mice per group). Data are expressed as mean ± s.e.m. Means between groups was compared with Student *t*-test. Statistical significance levels were set as follows: ** if p < 0.01, and *** if p < 0.001. ns: not statistically significant. PMN: neutrophils; DCs: dendritic cells.

**Fig. 3.**
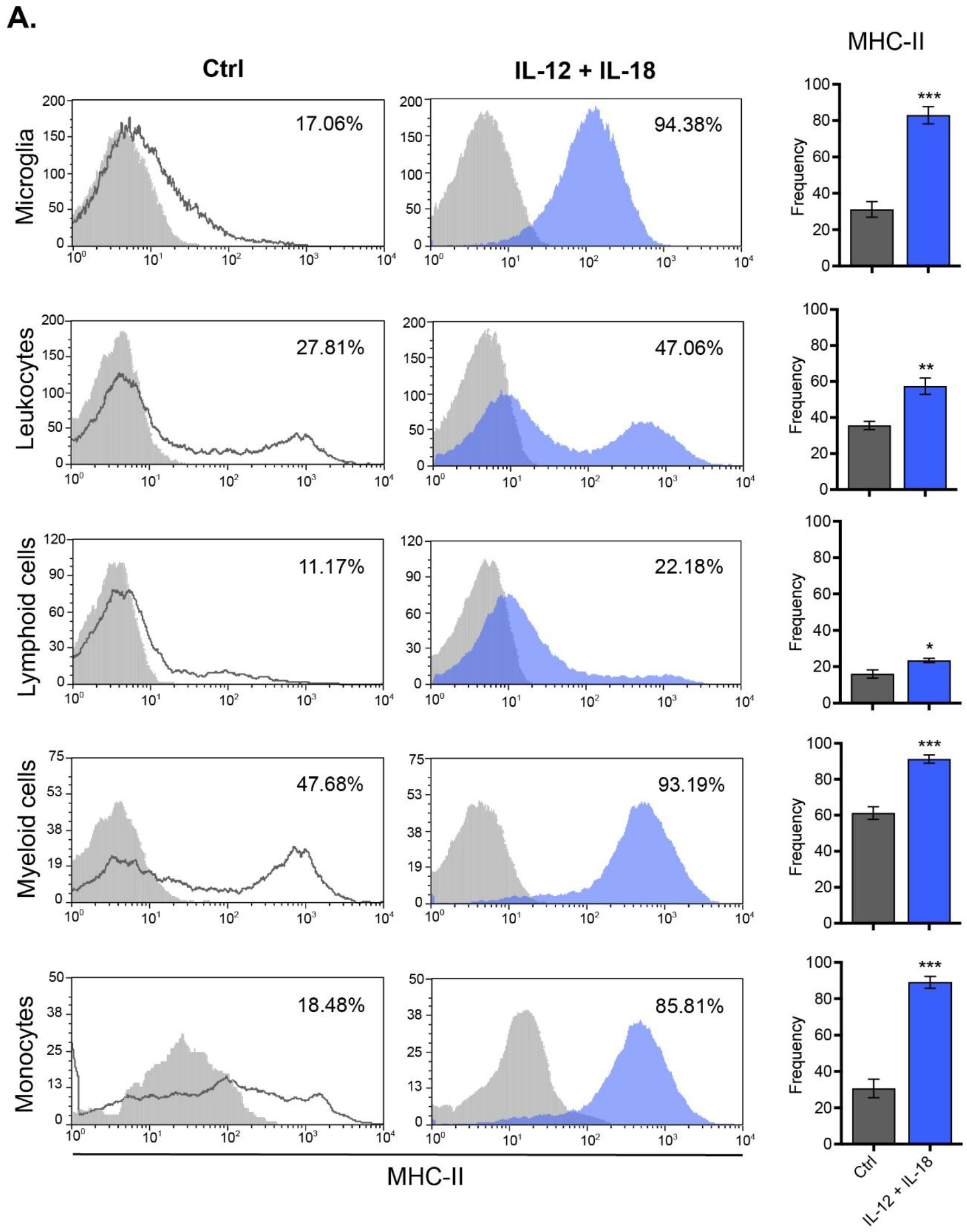
Peripheral delivery of IL-12 and IL-18 cDNAs leads to microglial activation and recruitment of MHC-II-expressing leukocytes toward the brain. C57BL/6 WT mice were hydrodynamically injected with either empty vector control cDNA (Ctrl), or IL-12 plus IL-18 cDNAs (IL-12 + IL-18). Seven days after initiation of the treatment mice were euthanized, and immune cells were isolated from whole brain and stained for subsequent flow cytometric analysis. **(A)** Representative stacked histograms depicting MHC-II expression across different immune cell populations. Frequencies of CD11b^+^ CD45^lo^ MHC-II^+^ microglia, CD11b^+/-^ CD45^hi^ MHC-II^+^ recruited leukocytes, CD11b^-^ CD45^hi^ MHC-II^+^ lymphoid cells, CD11b^+^ CD45^hi^ MHC-II^+^ myeloid cells, and CD11b^+^ CD45^hi^ Ly6C^hi^ Ly6G^-^ MHC-II^+^ inflammatory monocytes. Frequencies of all recruited immune cell subsets mentioned above, when gated in CD11b^+/-^ CD45^lo/hi^. Results are representative of at least three independent experiments (n= 4 mice per group). Data are expressed as mean ± s.e.m. Means between groups was compared with Student *t*-test. Statistical significance levels were set as follows: * if p < 0.05, ** if p < 0.01, and *** if p < 0.001.

Next, we found that IL-12 and IL-18 cDNAs-mediated neuroinflammation promoted the trafficking of CD45^hi^ leukocytes from the periphery and into the brain, resulting in a remodeling of the brain immune landscape (**Fig. 2E**). Interestingly, and in accordance with the heightened mRNA expression of CCR2 in the brain, we noticed a major contribution of CD45^hi^ CD11b^+^ Ly6C^hi^ inflammatory monocytes (**Fig. 2C**) and CD45^hi^ CD3^+^ CD8^+^ T cells (**Fig. 2D**) from the peripheral leukocyte compartment toward the brain. Similarly to what was shown for microglia, IL-12 plus IL-18-primed inflammatory monocytes did also upregulate MHC-II, relative to monocytes derived from empty vector-treated animals (**Fig. 3**).

In order to validate our previous findings, we next proceeded to assess microglial activation state and the specific localization of the leukocytes recruited into different brain areas by immunofluorescence, exploiting both the morphology and the dim expression of CD45 in star-shaped brain-resident microglia to distinguish them from rounded peripheral leukocytes bearing a bright expression of this same marker.

Following seven days after systemic expression of IL-12 plus IL-18 cDNAs, CD45^dim^ microglia effectively underwent activation, as determined by their integrated density area (**Fig. 4A and 4B**). This event led, in turn, to morphological alterations characterized by shortening and thickening of their processes along with an increase in the size of their cellular soma (**Fig. 4A**). Yet, we found no changes in microglial cell count per field, all results showing consistency with our previous flow cytometric analysis (**Fig. 4B**).

**Fig. 4.**
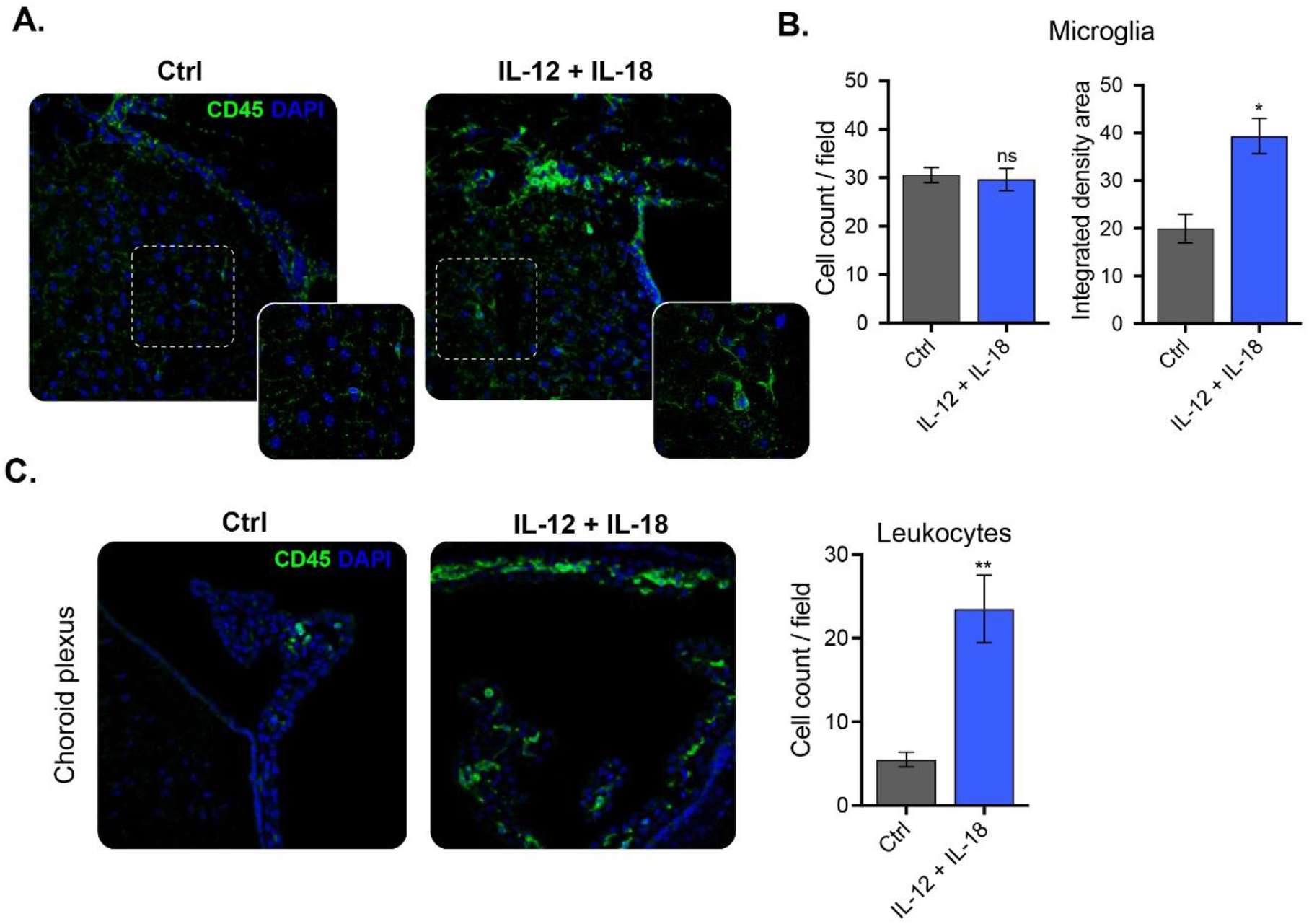
Peripheral challenge of IL-12 plus IL-18 induces leukocyte recruitment into the brain, through the choroid plexus. C57BL/6 WT mice were hydrodynamically injected with either empty vector control cDNA (Ctrl), or IL-12 plus IL-18 cDNAs (IL-12 + IL-18). Seven days after initiation of the treatment mice were euthanized, and brains collected. **(A)** Confocal micrographs depicting CD45^dim^ microglia (green, ramified cells), and CD45^bright^ recruited leukocytes (green, rounded cells). DAPI counterstain (blue) shows nucleus. Scale bars, 40 μm (main panels), 13 μm (inset). **(B)** Cell count per field and integrated density area of CD45^dim^ microglia (green, ramified cells). Confocal micrographs of **(C)** choroid plexus depicting CD45^dim^ microglia (green, ramified cells), CD45^bright^ recruited leukocytes (green, rounded cells). DAPI counterstain (blue) shows nucleus. Scale bars, 40 μm (main panels), 13 μm (inset). Cell count per field of CD45^bright^ recruited leukocytes (green, rounded cells). Results are representative of at least three independent experiments (n= 3-9 mice per group). Data are expressed as mean ± s.e.m. Means between groups was compared with Student *t*-test. Statistical significance levels were set as follows: * if p < 0.05, and ** if p < 0.01. ns: not statistically significant.

As for the CD45^bright^ leukocyte recruitment, we visualized a meaningful increase at the meninges and to a lesser extent, but still significant, into the caudate-putamen, greatly associated with vascular PECAM-expressing endothelial cells (**Fig. S1**). However, peripheral immune recruitment became substantially more evident at the choroid plexus (**Fig. 4C**).

In the present study, we propose a novel cytokine-mediated model of neuroinflammation through systemic delivery of IL-12 and IL-18 endotoxin-free cDNAs, to elucidate the impact of peripheral inflammation in leukocyte trafficking into the CNS. Since it has been reported in several experimental models that even minimal concentrations of LPS can activate the cerebral endothelium (2,12), and considering that we use bacteria as a means to amplify IL-12 plus IL-18-producing plasmids, we further validated our results in TLR4^-/-^ mice. Evaluation of splenomegaly (**Fig. S2A**), microglial and peripheral immune cell count (**Figs. S2B and S2C)** as well as their activation status (MHC-II expression) (**Fig. S3)**, showed no differences in all assessed parameters, between WT and TLR4 knock-out (KO) animals.

Taken together, these results clearly demonstrate that the systemic hydrodynamic administration of IL-12 and IL-18 cDNAs could be considered as an innovative experimental model of aseptic neuroinflammation, as it recapitulates all major features associated with this process in an endotoxin-free manner.

### IFN-γ is required to elicit peripheral inflammation, followed by microglial activation and leukocyte recruitment into the brain after systemic delivery of IL-12 plus IL-18 cDNAs

Given that TNF-α and IFN-γ were significantly induced both in the periphery and in the brain itself after systemic hydrodynamic shear of IL-12 and IL-18 cDNAs, we decided to address their involvement during neuroinflammation, by making use of TNF-αR1^-/-^ and IFN-γ^-/-^ mice. Therefore, we determined the circulating levels of cytokines, assessed splenomegaly, microglial activation, in addition to evaluation of peripheral immune cell trafficking into the brain in both mouse strains.

Following systemic expression of IL-12 plus IL-18, IFN-γ^-/-^ mice showed significantly low sera level of TNF-α and complete abrogation of IFN-γ, whereas TNF-αR1^-/-^ animals presented a kinetic and levels of TNF-α and IFN-γ comparable to their WT counterparts **(Fig. 5A)**. Moreover, since we did not notice signs of splenomegaly in IFN-γ^-/-^ mice **(Fig. 5B)**, we further validated our results by testing IL-12 protein level at different time points after cDNA delivery. Similar concentration of circulating IL-12 was dosed in IFN-γ^-/-^ mice in comparison to their WT littermates in all time points assessed, ruling out that the observed differences between transgenic strains were due to a less efficient hydrodynamic injection (**Fig. 5C**).

**Fig. 5.**
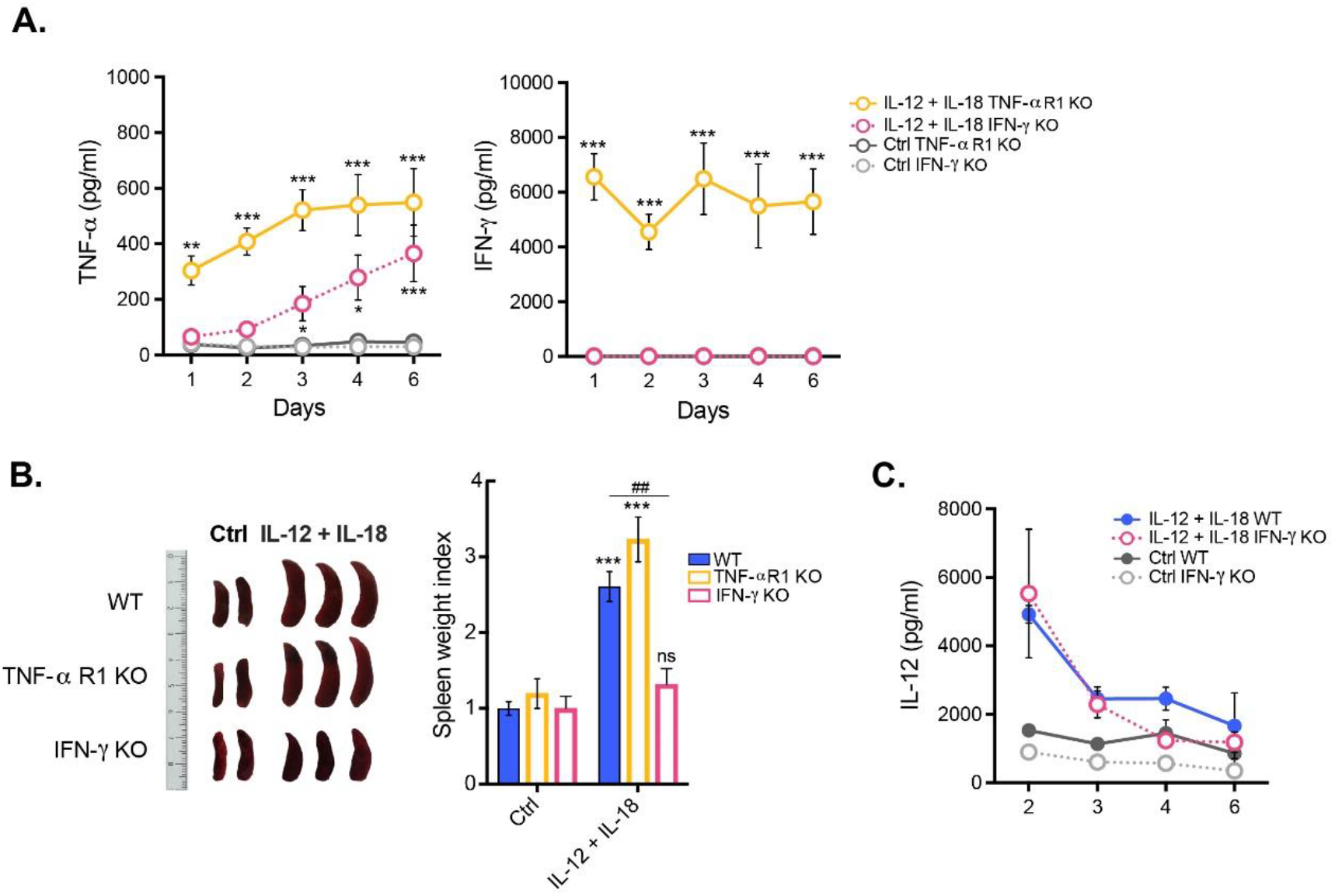
IFN-γ is required to exert peripheral inflammation after systemic induced-coexpression of IL-12 and IL-18. C57BL/6 WT, TNFαR1^-/-^ and IFN-γ^-/-^ mice were hydrodynamically injected with either empty vector control cDNA (Ctrl), or IL-12 plus IL-18 cDNAs (IL-12 + IL-18). **(A)** Serum samples obtained at day 0 and at different time points following initiation of the treatment were tested for TNF-α and IFN-γ. Results are representative of at least three independent experiments (n= 3-7 samples per time point and group). **(B)** Seven days after initiation of the treatment mice were euthanized, and spleens excised for assessment of their relative weight and size. Results are representative of at least three independent experiments (n= 2-4 mice per group). **(C)** Serum samples obtained at day 0 and at different time points following initiation of the treatment were tested for IL-12. Results are representative of at least three independent experiments (n= 3-4 mice per group). Data are expressed as mean ± s.e.m. Means between groups was compared with two-way analysis of variance followed by a Bonferroni *post-hoc* test. Statistical significance levels were set as follows: * if p < 0.05, **/## if p < 0.01, and *** if p < 0.001. ns: not statistically significant.

The absence of splenomegaly in IFN-γ KO prompted us to test whether IFN-γ could be necessary and sufficient to mount a systemic cytokine-mediated two-step process after delivery of IL-12 plus IL-18 cDNAs, initiating an enhanced peripheral inflammation, which could in turn lead to neuroinflammation, featured by activation of microglia and recruitment of leukocytes into the brain. Notably, we observed that only IFN-γ, but not TNF-α signaling through its type I receptor, was required to both induce the trafficking of leukocytes from the periphery toward the brain (**Fig. 6**) and upregulate MHC-II in microglia and inflammatory monocytes **(Fig. 7)**.

**Fig. 6.**
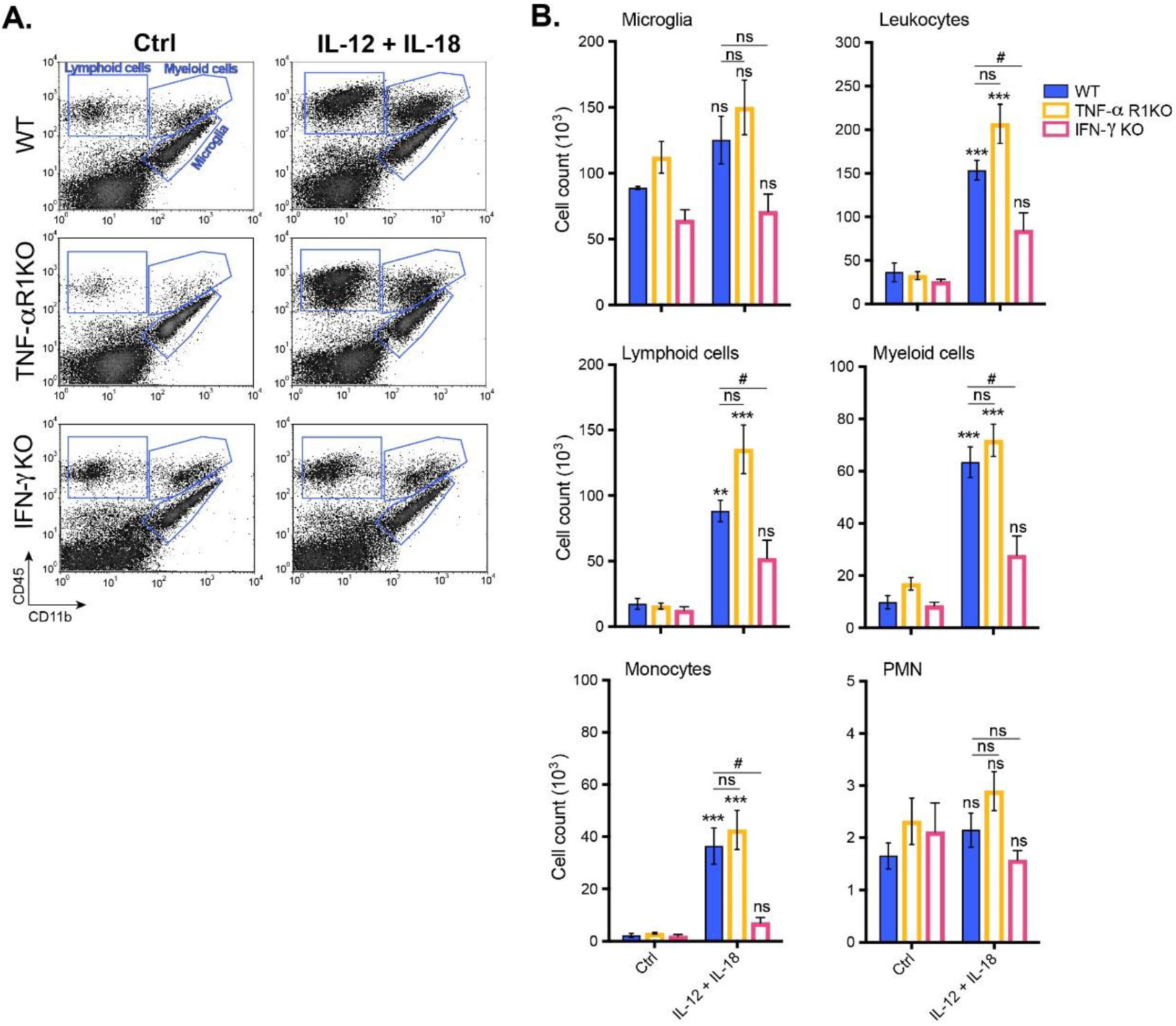
IFN-γ deficiency results in impaired leukocyte recruitment toward the brain upon systemic delivery of IL-12 and IL-18 cDNAs. C57BL/6 WT, TNFαR1^-/-^ and IFN-γ^-/-^ mice were hydrodynamically injected with either empty vector control cDNA (Ctrl), or IL-12 plus IL-18 cDNAs (IL-12 + IL-18). Seven days after initiation of the treatment mice were euthanized, and immune cells were isolated from whole brain and stained for subsequent flow cytometric analysis. **(A)** Representative CD45 vs. CD11b flow cytometric density-plots illustrating the gating analysis strategy employed. **(B)** Absolute number of CD11b^+^ CD45^lo^ microglia, CD11b^+/-^ CD45^hi^ recruited leukocytes, CD11b^-^ CD45^hi^ recruited lymphoid cells, CD11b^+^ CD45^hi^ recruited myeloid cells, CD11b^+^ CD45^hi^ Ly6C^hi^ inflammatory monocytes, and CD11b^+^ CD45^hi^ Ly6G^hi^ neutrophils were assessed by flow cytometry. Results are representative of at least three independent experiments (n= 2-7 mice per group). Data are expressed as mean ± s.e.m. Means between groups was compared with two-way analysis of variance followed by a Bonferroni *post-hoc* test. Statistical significance levels were set as follows: # if p < 0.05, ** if p < 0.01, and *** if p < 0.001. ns: not statistically significant.

**Fig. 7.**
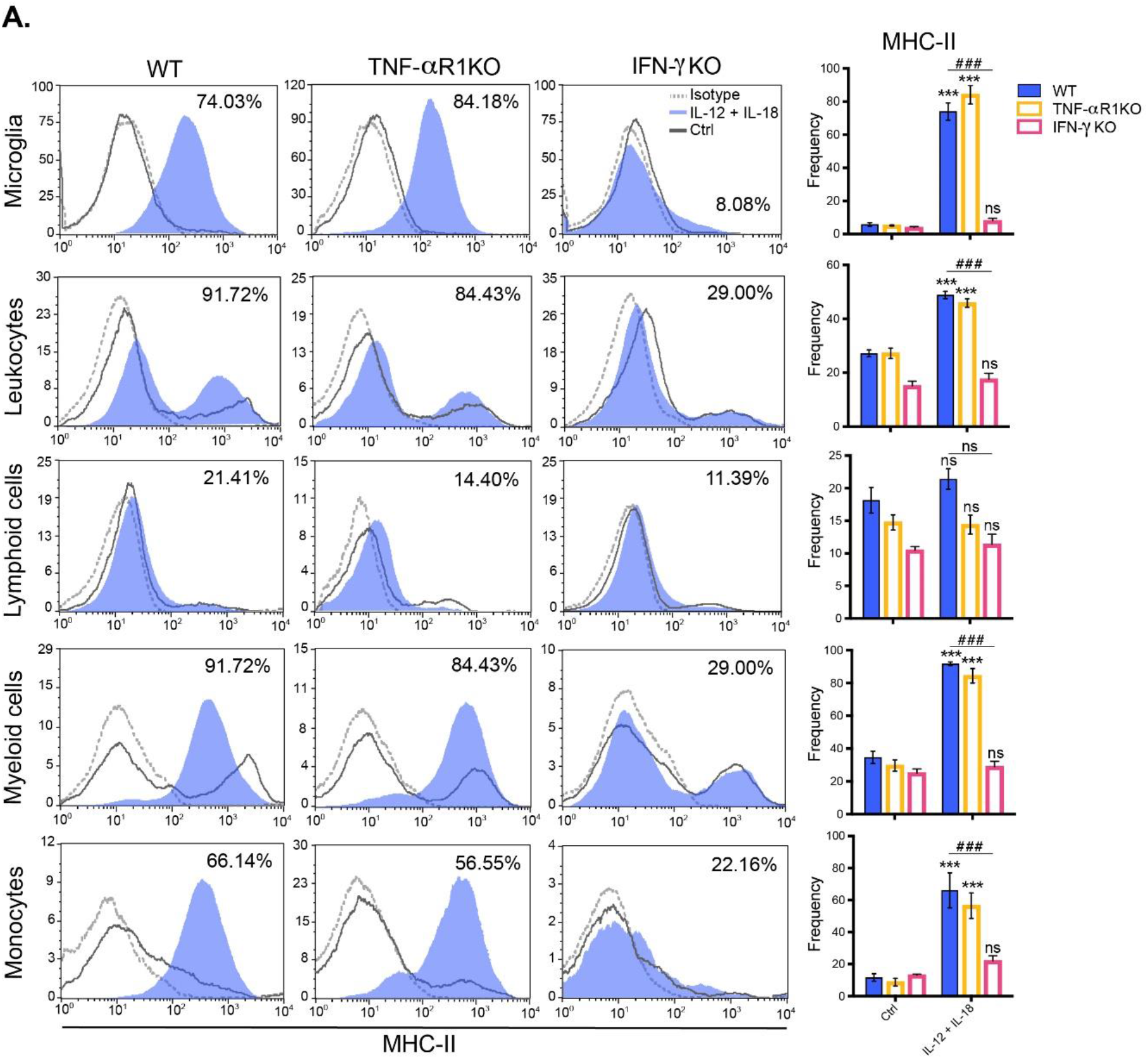
Systemic hydrodynamic shear of IL-12 plus IL-18 cDNAs drives IFN-γ-dependent activation of microglia and recruitment of MHC-II-expressing monocytes into the brain. C57BL/6 WT, TNFαR1^-/-^ and IFN-γ^-/-^ mice were hydrodynamically injected with either empty vector control cDNA (Ctrl), or IL-12 plus IL-18 cDNAs (IL-12 + IL-18). Seven days after initiation of the treatment mice were euthanized, and immune cells were isolated from whole brain and stained for subsequent flow cytometric analysis. **(A)** Representative stacked histograms depicting MHC-II expression across different immune cell populations. Frequencies of CD11b^+^ CD45^lo^ MHC-II^+^ microglia, CD11b^+/-^ CD45^hi^ MHC-II^+^ recruited leukocytes, CD11b^-^ CD45^hi^ MHC-II^+^ recruited lymphoid cells, CD11b^+^ CD45^hi^ MHC-II^+^ recruited myeloid cells, and CD11b^+^ CD45^hi^ Ly6C^hi^ MHC-II^+^ inflammatory monocytes. Results are representative of at least three independent experiments (n= 4-7 mice per group). Data are expressed as mean ± s.e.m. Means between groups was compared with two-way analysis of variance followed by a Bonferroni *post-hoc* test. Statistical significance levels were set as follows: ^***/###^ if p < 0.001. ns: not statistically significant.

Overall, our results point to an IFN-γ-dependent immune response triggered by systemic sterile induced-co-expression of IL-12 and IL-18, which leads to activation of microglia and leukocyte recruitment into the brain during the onset and development of this cytokine-mediated model of neuroinflammation.

## DISCUSSION

Here we report the novel effect of systemic, sterile induced-co-expression of IL-12 and IL-18 in the establishment of an innovative cytokine-mediated murine model of neuroinflammation, through peripheral hydrodynamic shear of cDNA.

Our results show that peripheral inflammation is mediated by a cytokine storm including persistent high levels of IL-12, TNF-α and IFN-γ, along with a pronounced splenomegaly, which thereof became a distinctive trait, and more important, a control of an efficient treatment. Accordingly, it has been previously shown that an increase in IL-10 correlates with a protective role mediated by this cytokine, limiting an exacerbated inflammation induced by pro-inflammatory cytokines such as IL-12 (26) and IFN-γ (39). Besides, TNF-α was associated with hepatic toxicity and diminished mice survival when IL-12 cDNA was injected alone (32). Thus, this coordinated Th1/Th2 balance is critical to shape a well-controlled peripheral immune response against systemic inflammatory events, which could in turn lead to neuroinflammation.

Herein, we demonstrate that this cytokine-mediated process unleash the peripheral immune system, inducing the mobilization of highly activated inflammatory monocytes and CD8^+^ T cells into the brain, harnessing MHC-II^+^ tissue-resident microglia in an IFN-γ-mediated manner (40), suggesting that microglia interaction with Th1 cells would license them with potential antigen-presenting function, as reported before (41). Moreover, we found an augment in the mRNA gene expression of CCR2 in the brain, which has been widely described as a key chemokine receptor orchestrating monocyte (42) and T cell (43) migration to inflamed sites. Interestingly, we localized significantly more immune cells at the choroid plexus, which constitute an epithelial layer of the blood cerebrospinal-fluid barrier that has been postulated to be a unique IFN-γ-dependent neuroimmunological interface between the periphery and the brain, displaying a key role as a gateway for CNS immune surveillance (37,44,45). This solely would explain the impairment in the activation and recruitment of leukocytes into the brain of IFN-γ KO mice after systemic delivery of IL-12 and IL-18 cDNAs, supporting our hypothesis of neuroinflammation as a two-step process that may start in the periphery and finalizes locally in the brain. However, it remains to be elucidated whether IFN-γ peripherally or locally expressed at the choroid plexus would be the responsible for modulating leukocyte trafficking into the brain.

Collectively, the results presented argue for a systemic cytokine-mediated origin of successive immunological events for the establishment and development of neuroinflammation, having identified IFN-γ as a potential target for immunotherapy.

## MATERIALS AND DATA AVAILABILITY STATEMENT

The datasets used and/or analyzed during the current study are available from the corresponding author on reasonable request.

## AUTHOR CONTRIBUTIONS

EAG conceived and designed the research study, performed the experiments, analyzed data and wrote the manuscript. JMPR performed the experiments, aided in interpreting the results, prepare the data for presentation and wrote the manuscript. DSA performed the experiments and together with CB discussed and commented on the manuscript. PI conceived and designed the research study, aided in interpreting the results, and supervised the manuscript. MCRG designed the research study, aided in interpreting the results, and wrote the manuscript. All authors reviewed the manuscript before submission.

## ACKNOWLEDGEMENTS

The authors thank Fabricio Navarro, Diego Luti, Carolina Florit, Victoria Blanco, and Ivanna Novotny-Núñez for animal care, Pilar Crespo and Paula Abadie for FACS technical support and Paula Icely for overall experimental technical assistance. Authors would especially like to thank Dr. Denise Kviatcovsky for kindly designing the figures and for insightful discussion on the manuscript.

## SUPPLEMENTARY FIGURES

**Fig. S1.**
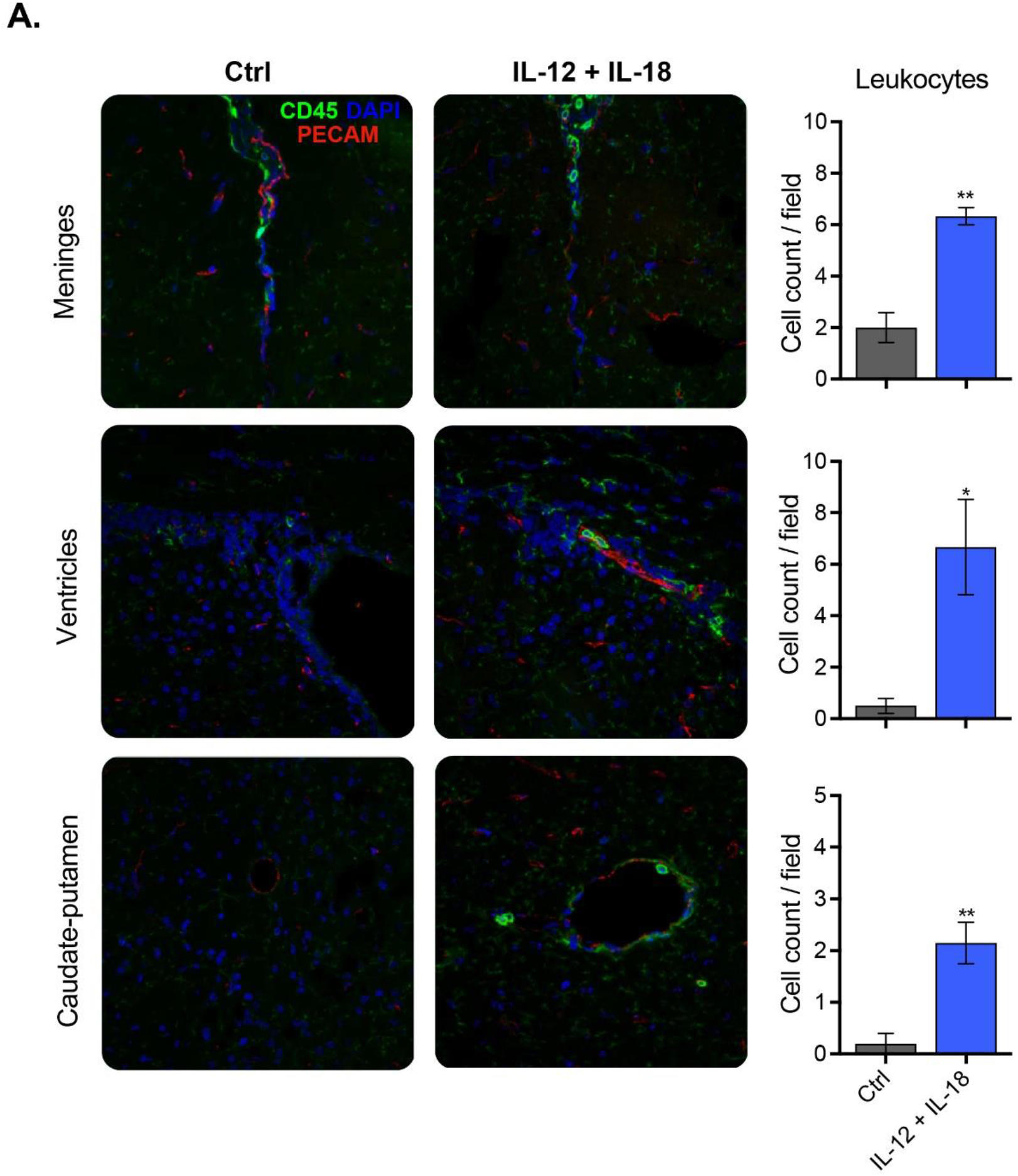
Peripheral challenge of IL-12 plus IL-18 cDNAs leads to leukocyte recruitment into distinct brain regions. C57BL/6 WT mice were hydrodynamically injected with either empty vector control cDNA (Ctrl) or IL-12 plus IL-18 cDNAs (IL-12 + IL-18). Seven days after initiation of the treatment mice were euthanized, and brains collected. Confocal micrographs of meninges, ventricles, and caudate-putamen depicting CD45^dim^ microglia (green, ramified cells), CD45^bright^ recruited leukocytes (green, rounded cells), and PECAM^bright^ endothelial cells (red). DAPI counterstain (blue) shows nucleus. Scale bars, 40 μm. Cell count per field of CD45^bright^ recruited leukocytes (green, rounded cells). Results are representative of at least three independent experiments (n= 3-7 mice per group). Data are expressed as mean ± s.e.m. Means between groups was compared with Student *t*-test. Statistical significance levels were set as follows: * if p < 0.05, and ** if p < 0.01.

**Fig. S2.**
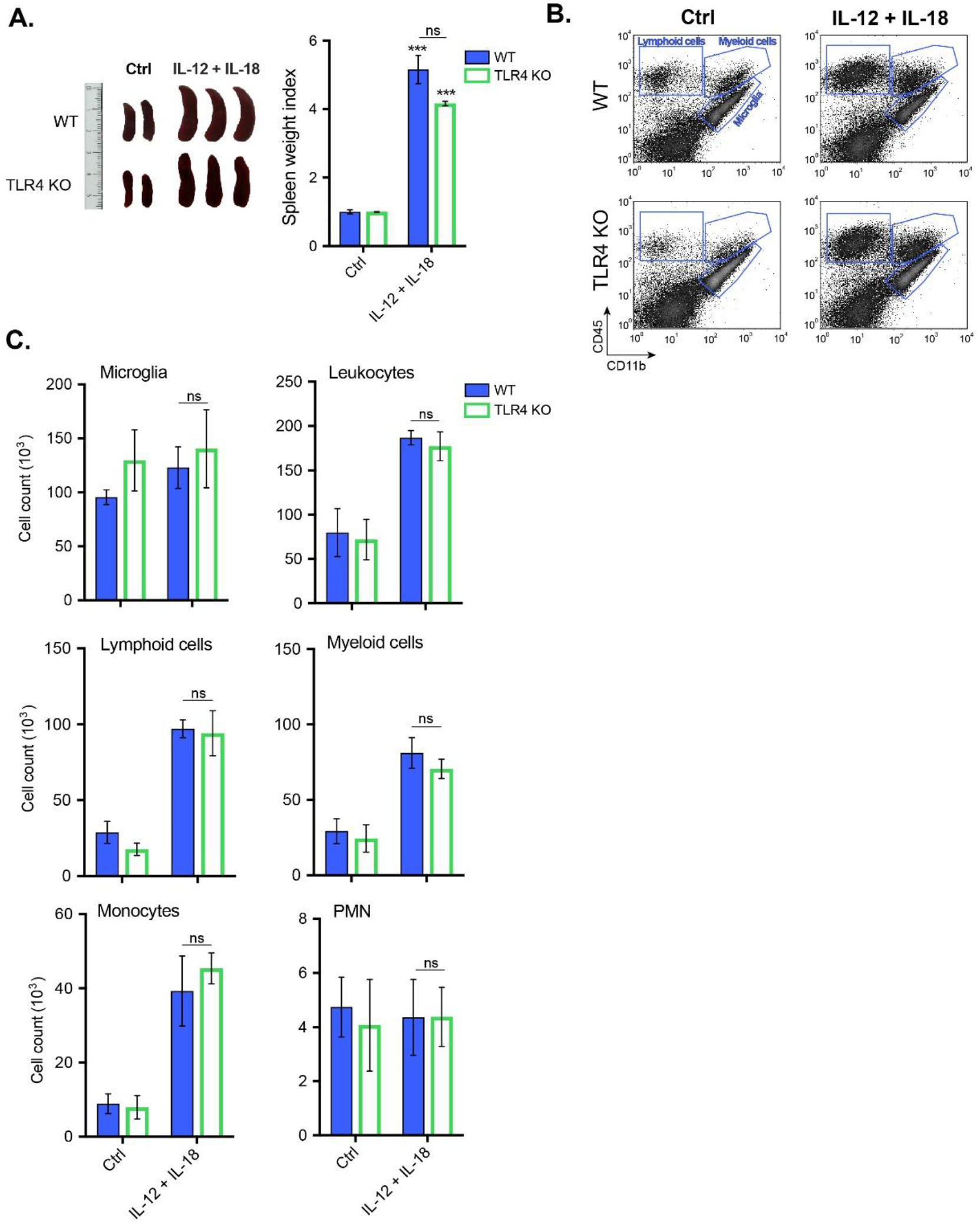
TLR4 is not required to induce peripheral inflammation after systemic induced-co-expression of IL-12 and IL-18. C57BL/6 WT, and TLR4^-/-^ mice were hydrodynamically injected with either empty vector control cDNA (Ctrl) or IL-12 plus IL-18 cDNAs (IL-12 + IL-18). **(A)** Seven days after initiation of the treatment mice were euthanized, and spleens excised for assessment of their relative weight and size. Results are representative of at least three independent experiments (n= 2-3 mice per group). **(B)** Immune cells were isolated from whole brain and stained for subsequent flow cytometric analysis. Representative CD45 vs. CD11b flow cytometric density-plots illustrating the gating analysis strategy employed. **(C)** Absolute number of CD11b^+^ CD45^lo^ microglia, CD11b^+/-^ CD45^hi^ recruited leukocytes, CD11b^+^ CD45^hi^ recruited myeloid cells, CD11b^-^ CD45^hi^ recruited lymphoid cells, CD11b^+^ CD45^hi^ Ly6C^hi^ inflammatory monocytes, and CD11b^+^ CD45^hi^ Ly6G^hi^ neutrophils were assessed by flow cytometry. Results are representative of at least three independent experiments (n= 2-4 mice per group). Data are expressed as mean ± s.e.m. Means between groups was compared with two-way analysis of variance followed by a Bonferroni *post-hoc* test. Statistical significance levels were set as follows: *** if p < 0.001. ns: not statistically significant.

**Fig. S3.**
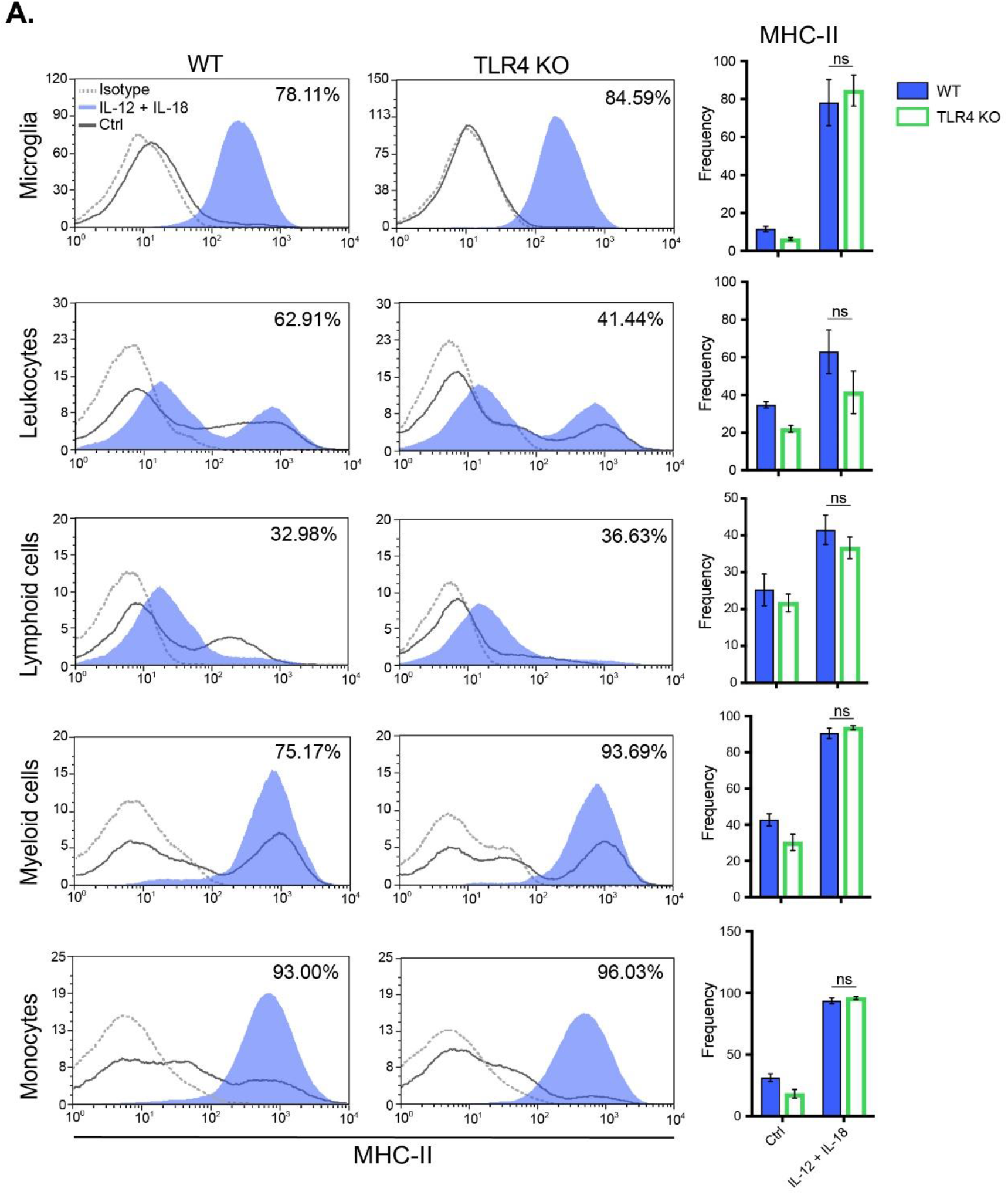
Peripheral sterile hydrodynamic shear of IL-12 plus IL-18 cDNAs induces activation of microglia and recruitment of MHC-II-expressing monocytes into the brain in a TLR4-independent manner. C57BL/6 WT, and TLR4^-/-^ mice were hydrodynamically injected with either empty vector control cDNA (Ctrl) or IL-12 plus IL-18 cDNAs (IL-12 + IL-18). Seven days after initiation of the treatment mice were euthanized, and immune cells were isolated from whole brain and stained for subsequent flow cytometric analysis. **(A)** Representative stacked histograms depicting MHC-II expression. **(B)** Frequencies of CD11b^+^ CD45^lo^ MHC-II^+^ microglia, CD11b^+/-^ CD45^hi^ MHC-II^+^ recruited leukocytes, CD11b^-^ CD45^hi^ MHC-II^+^ recruited lymphoid cells, CD11b^+^ CD45^hi^ MHC-II^+^ recruited myeloid cells, and CD11b^+^ CD45^hi^ Ly6C^hi^ MHC-II^+^ inflammatory monocytes. Results are representative of at least three independent experiments (n= 2-4 mice per group). Data are expressed as mean ± s.e.m. Means between groups was compared with two-way analysis of variance followed by a Bonferroni *post-hoc* test. ns: not statistically significant.

## REFERENCES

1. Perry VH, Newman TA, Cunningham C. The impact of systemic infection on the progression of neurodegenerative disease. Nat Rev Neurosci (2003) 4:103–112. doi:10.1038/nrn1032

2. Cunningham C. Microglia and neurodegeneration: the role of systemic inflammation. Glia (2013) 61:71–90. doi:10.1002/glia.22350

3. Neher JJ, Cunningham C. Priming microglia for innate immune memory in the brain. Trends Immunol (2019) 40:358–374. doi:10.1016/j.it.2019.02.001

4. Wendeln A-C, Degenhardt K, Kaurani L, Gertig M, Ulas T, Jain G, Wagner J, Häsler LM, Wild K, Skodras A, et al. Innate immune memory in the brain shapes neurological disease hallmarks. Nature (2018) 556:332–338. doi:10.1038/s41586-018-0023-4

5. Arroyo DS, Gaviglio EA, Peralta Ramos JM, Bussi C, Avalos MP, Cancela LM, Iribarren P. Phosphatidyl-Inositol-3 Kinase Inhibitors Regulate Peptidoglycan-Induced Myeloid Leukocyte Recruitment, Inflammation, and Neurotoxicity in Mouse Brain. Front Immunol (2018) 9:770. doi:10.3389/fimmu.2018.00770

6. Rua R, Lee JY, Silva AB, Swafford IS, Maric D, Johnson KR, McGavern DB. Infection drives meningeal engraftment by inflammatory monocytes that impairs CNS immunity. Nat Immunol (2019) 20:407–419. doi:10.1038/s41590-019-0344-y

7. Winkler CW, Woods TA, Robertson SJ, McNally KL, Carmody AB, Best SM, Peterson KE. Cutting Edge: CCR2 Is Not Required for Ly6Chi Monocyte Egress from the Bone Marrow but Is Necessary for Migration within the Brain in La Crosse Virus Encephalitis. J Immunol (2018) 200:471–476. doi:10.4049/jimmunol.1701230

8. Wang A, Huen SC, Luan HH, Baker K, Rinder H, Booth CJ, Medzhitov R. Glucose metabolism mediates disease tolerance in cerebral malaria. Proc Natl Acad Sci USA (2018) 115:11042–11047. doi:10.1073/pnas.1806376115

9. Mahamed DA, Mills JH, Egan CE, Denkers EY, Bynoe MS. CD73-generated adenosine facilitates Toxoplasma gondii differentiation to long-lived tissue cysts in the central nervous system. Proc Natl Acad Sci USA (2012) 109:16312–16317. doi:10.1073/pnas.1205589109

10. Zhou H, Lapointe BM, Clark SR, Zbytnuik L, Kubes P. A requirement for microglial TLR4 in leukocyte recruitment into brain in response to lipopolysaccharide. J Immunol (2006) 177:8103–8110. doi:10.4049/jimmunol.177.11.8103

11. Cardona AE, Pioro EP, Sasse ME, Kostenko V, Cardona SM, Dijkstra IM, Huang D, Kidd G, Dombrowski S, Dutta R, et al. Control of microglial neurotoxicity by the fractalkine receptor. Nat Neurosci (2006) 9:917–924. doi:10.1038/nn1715

12. Hoogland ICM, Houbolt C, van Westerloo DJ, van Gool WA, van de Beek D. Systemic inflammation and microglial activation: systematic review of animal experiments. J Neuroinflammation (2015) 12:114. doi:10.1186/s12974-015-0332-6

13. Peralta Ramos JM, Bussi C, Gaviglio EA, Arroyo DS, Baez NS, Rodriguez-Galan MC, Iribarren P. Type I IFNs Are Required to Promote Central Nervous System Immune Surveillance through the Recruitment of Inflammatory Monocytes upon Systemic Inflammation. Front Immunol (2017) 8:1666. doi:10.3389/fimmu.2017.01666

14. Peralta Ramos JM, Iribarren P, Bousset L, Melki R, Baekelandt V, Van der Perren A. Peripheral Inflammation Regulates CNS Immune Surveillance Through the Recruitment of Inflammatory Monocytes Upon Systemic α-Synuclein Administration. Front Immunol (2019) 10:80. doi:10.3389/fimmu.2019.00080

15. Mazzolini G, Prieto J, Melero I. Gene therapy of cancer with interleukin-12. Curr Pharm Des (2003) 9:1981–1991. doi:10.2174/1381612033454261

16. Rakhmilevich AL, Timmins JG, Janssen K, Pohlmann EL, Sheehy MJ, Yang NS. Gene gun-mediated IL-12 gene therapy induces antitumor effects in the absence of toxicity: a direct comparison with systemic IL-12 protein therapy. J Immunother (1999) 22:135–144. doi:10.1097/00002371-199903000-00005

17. Rook AH, Zaki MH, Wysocka M, Wood GS, Duvic M, Showe LC, Foss F, Shapiro M, Kuzel TM, Olsen EA, et al. The role for interleukin-12 therapy of cutaneous T cell lymphoma. Ann N Y Acad Sci (2001) 941:177–184. doi:10.1111/j.1749-6632.2001.tb03721.x

18. Shurin MR, Esche C, Péron JM, Lotze MT. Antitumor activities of IL-12 and mechanisms of action. Chem Immunol (1997) 68:153–174.

19. Chiocca EA, Yu JS, Lukas RV, Solomon IH, Ligon KL, Nakashima H, Triggs DA, Reardon DA, Wen P, Stopa BM, et al. Regulatable interleukin-12 gene therapy in patients with recurrent high-grade glioma: Results of a phase 1 trial. Sci Transl Med (2019) 11: doi:10.1126/scitranslmed.aaw5680

20. Cua DJ, Sherlock J, Chen Y, Murphy CA, Joyce B, Seymour B, Lucian L, To W, Kwan S, Churakova T, et al. Interleukin-23 rather than interleukin-12 is the critical cytokine for autoimmune inflammation of the brain. Nature (2003) 421:744–748. doi:10.1038/nature01355

21. Eede P, Obst J, Benke E, Yvon-Durocher G, Richard BC, Gimber N, Schmoranzer J, Böddrich A, Wanker EE, Prokop S, et al. Interleukin-12/23 deficiency differentially affects pathology in male and female Alzheimer’s disease-like mice. EMBO Rep (2020) 21:e48530. doi:10.15252/embr.201948530

22. Vom Berg J, Prokop S, Miller KR, Obst J, Kälin RE, Lopategui-Cabezas I, Wegner A, Mair F, Schipke CG, Peters O, et al. Inhibition of IL-12/IL-23 signaling reduces Alzheimer’s disease-like pathology and cognitive decline. Nat Med (2012) 18:1812–1819. doi:10.1038/nm.2965

23. Lampa J, Westman M, Kadetoff D, Agréus AN, Le Maître E, Gillis-Haegerstrand C, Andersson M, Khademi M, Corr M, Christianson CA, et al. Peripheral inflammatory disease associated with centrally activated IL-1 system in humans and mice. Proc Natl Acad Sci USA (2012) 109:12728–12733. doi:10.1073/pnas.1118748109

24. Jha S, Srivastava SY, Brickey WJ, Iocca H, Toews A, Morrison JP, Chen VS, Gris D, Matsushima GK, Ting JP-Y. The inflammasome sensor, NLRP3, regulates CNS inflammation and demyelination via caspase-1 and interleukin-18. J Neurosci (2010) 30:15811–15820. doi:10.1523/JNEUROSCI.4088-10.2010

25. Saresella M, Basilico N, Marventano I, Perego F, La Rosa F, Piancone F, Taramelli D, Banks H, Clerici M. Leishmania infantum infection reduces the amyloid β42-stimulated NLRP3 inflammasome activation. Brain Behav Immun (2020) doi:10.1016/j.bbi.2020.04.058

26. Rodriguez-Galan MC, Reynolds D, Correa SG, Iribarren P, Watanabe M, Young HA. Coexpression of IL-18 strongly attenuates IL-12-induced systemic toxicity through a rapid induction of IL-10 without affecting its antitumor capacity. J Immunol (2009) 183:740–748. doi: 10.4049/jimmunol.0804166

27. Rodriguez-Galán MC, Bream JH, Farr A, Young HA. Synergistic effect of IL-2, IL-12, and IL-18 on thymocyte apoptosis and Th1/Th2 cytokine expression. J Immunol (2005) 174:2796–2804. doi:10.4049/jimmunol.174.5.2796

28. Zhou T, Damsky W, Weizman O-E, McGeary MK, Hartmann KP, Rosen CE, Fischer S, Jackson R, Flavell RA, Wang J, et al. IL-18BP is a secreted immune checkpoint and barrier to IL-18 immunotherapy. Nature (2020) doi:10.1038/s41586-020-2422-6

29. Felderhoff-Mueser U, Schmidt OI, Oberholzer A, Bührer C, Stahel PF. IL-18: a key player in neuroinflammation and neurodegeneration? Trends Neurosci (2005) 28:487–493. doi: 10.1016/j.tins.2005.06.008

30. Voet S, Srinivasan S, Lamkanfi M, van Loo G. Inflammasomes in neuroinflammatory and neurodegenerative diseases. EMBO Mol Med (2019) 11: doi:10.15252/emmm.201810248

31. Liu F, Song Y, Liu D. Hydrodynamics-based transfection in animals by systemic administration of plasmid DNA. Gene Ther (1999) 6:1258–1266. doi:10.1038/sj.gt.3300947

32. Barrios B, Baez NS, Reynolds D, Iribarren P, Cejas H, Young HA, Rodriguez-Galan MC. Abrogation of TNFα production during cancer immunotherapy is crucial for suppressing side effects due to the systemic expression of IL-12. PLoS One (2014) 9:e90116. doi: 10.1371/journal.pone.0090116

33. Baez NS, Cerbán F, Savid-Frontera C, Hodge DL, Tosello J, Acosta-Rodriguez E, Almada L, Gruppi A, Viano ME, Young HA, et al. Thymic expression of IL-4 and IL-15 after systemic inflammatory or infectious Th1 disease processes induce the acquisition of “innate” characteristics during CD8+ T cell development. PLoS Pathog (2019) 15:e1007456. doi:10.1371/journal.ppat.1007456

34. Qiu D, Chu X, Hua L, Yang Y, Li K, Han Y, Yin J, Zhu M, Mu S, Sun Z, et al. Gpr174- deficient regulatory T cells decrease cytokine storm in septic mice. Cell Death Dis (2019) 10:233. doi:10.1038/s41419-019-1462-z

35. Chousterman BG, Swirski FK, Weber GF. Cytokine storm and sepsis disease pathogenesis. Semin Immunopathol (2017) 39:517–528. doi:10.1007/s00281-017-0639-8

36. Ousman SS, Kubes P. Immune surveillance in the central nervous system. Nat Neurosci (2012) 15:1096–1101. doi:10.1038/nn.3161

37. Schwartz M, Peralta Ramos JM, Ben-Yehuda H. A 20-Year Journey from Axonal Injury to Neurodegenerative Diseases and the Prospect of Immunotherapy for Combating Alzheimer’s Disease. J Immunol (2020) 204:243–250. doi:10.4049/jimmunol.1900844

38. Prinz M, Priller J. The role of peripheral immune cells in the CNS in steady state and disease. Nat Neurosci (2017) 20:136–144. doi:10.1038/nn.4475

39. Savarin C, Bergmann CC. Fine Tuning the Cytokine Storm by IFN and IL-10 Following Neurotropic Coronavirus Encephalomyelitis. Front Immunol (2018) 9:3022. doi:10.3389/fimmu.2018.03022

40. Panek RB, Benveniste EN. Class II MHC gene expression in microglia. Regulation by the cytokines IFN-gamma, TNF-alpha, and TGF-beta. J Immunol (1995) 154:2846–2854.

41. Aloisi F, De Simone R, Columba-Cabezas S, Penna G, Adorini L. Functional maturation of adult mouse resting microglia into an APC is promoted by granulocyte-macrophage colony-stimulating factor and interaction with Th1 cells. J Immunol (2000) 164:1705–1712. doi:10.4049/jimmunol.164.4.1705

42. Croxford AL, Lanzinger M, Hartmann FJ, Schreiner B, Mair F, Pelczar P, Clausen BE, Jung S, Greter M, Becher B. The Cytokine GM-CSF Drives the Inflammatory Signature of CCR2+ Monocytes and Licenses Autoimmunity. Immunity (2015) 43:502–514. doi:10.1016/j.immuni.2015.08.010

43. Rao DA, Gurish MF, Marshall JL, Slowikowski K, Fonseka CY, Liu Y, Donlin LT, Henderson LA, Wei K, Mizoguchi F, et al. Pathologically expanded peripheral T helper cell subset drives B cells in rheumatoid arthritis. Nature (2017) 542:110–114. doi: 10.1038/nature20810

44. Schwartz M, Baruch K. The resolution of neuroinflammation in neurodegeneration: leukocyte recruitment via the choroid plexus. EMBO J (2014) 33:7–22. doi: 10.1002/embj.201386609

45. Kunis G, Baruch K, Rosenzweig N, Kertser A, Miller O, Berkutzki T, Schwartz M. IFN-γ- dependent activation of the brain’s choroid plexus for CNS immune surveillance and repair. Brain (2013) 136:3427–3440. doi:10.1093/brain/awt259

